## Supplementary figures and images for "A non-canonical role for the autophagy machinery in anti-retroviral signaling mediated by TRIM5α"

### Figure S1

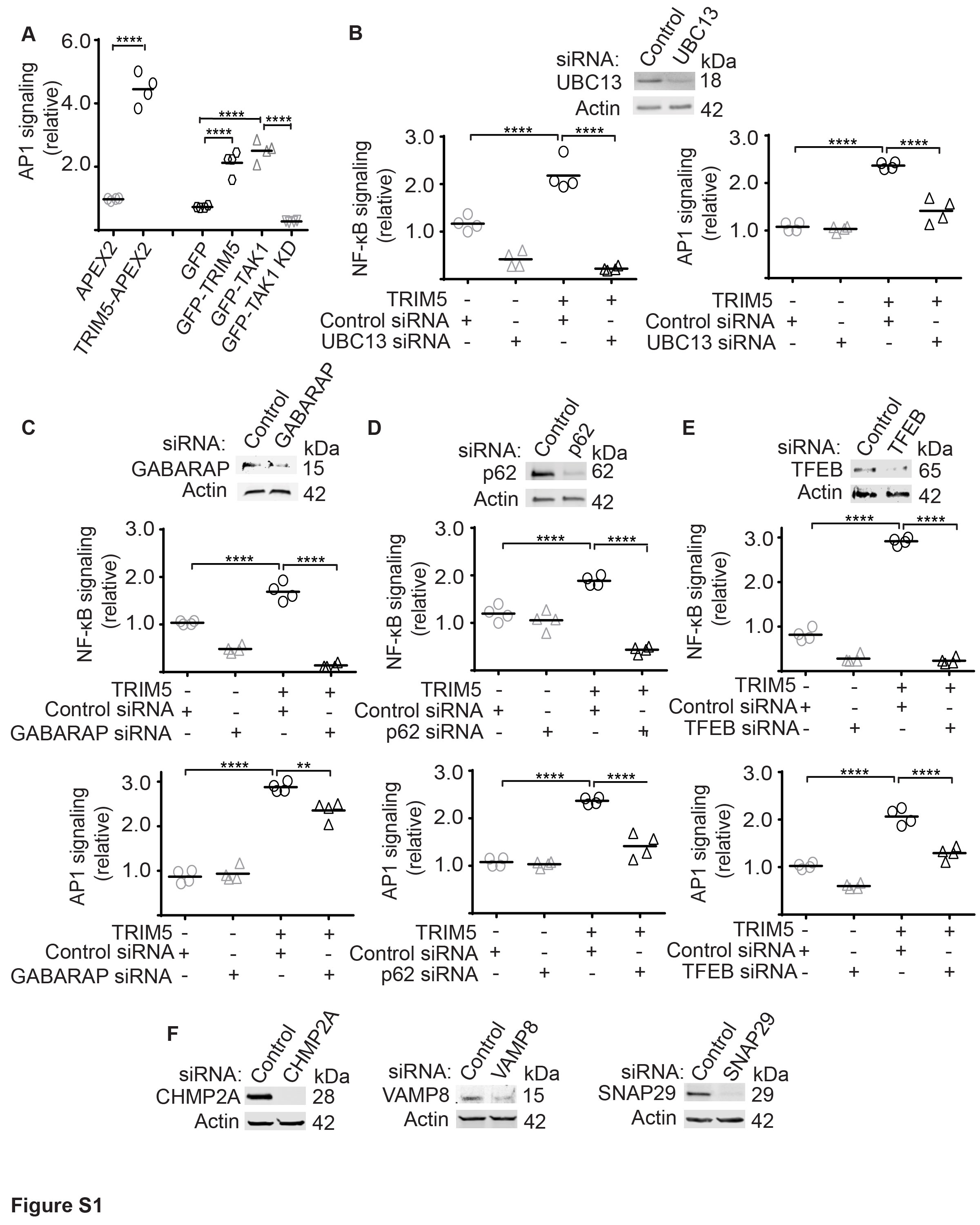

### Figure S2

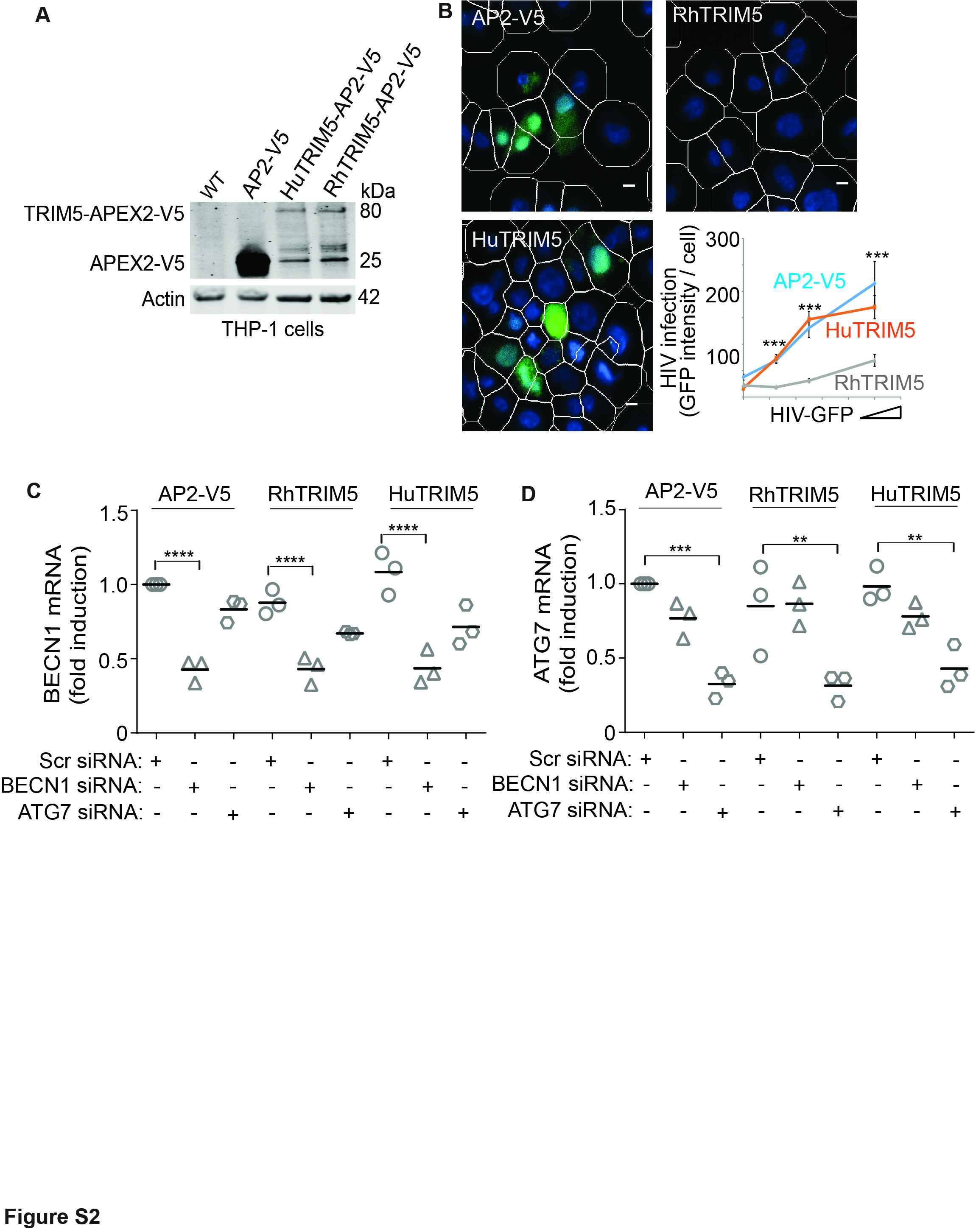

### Figure S3

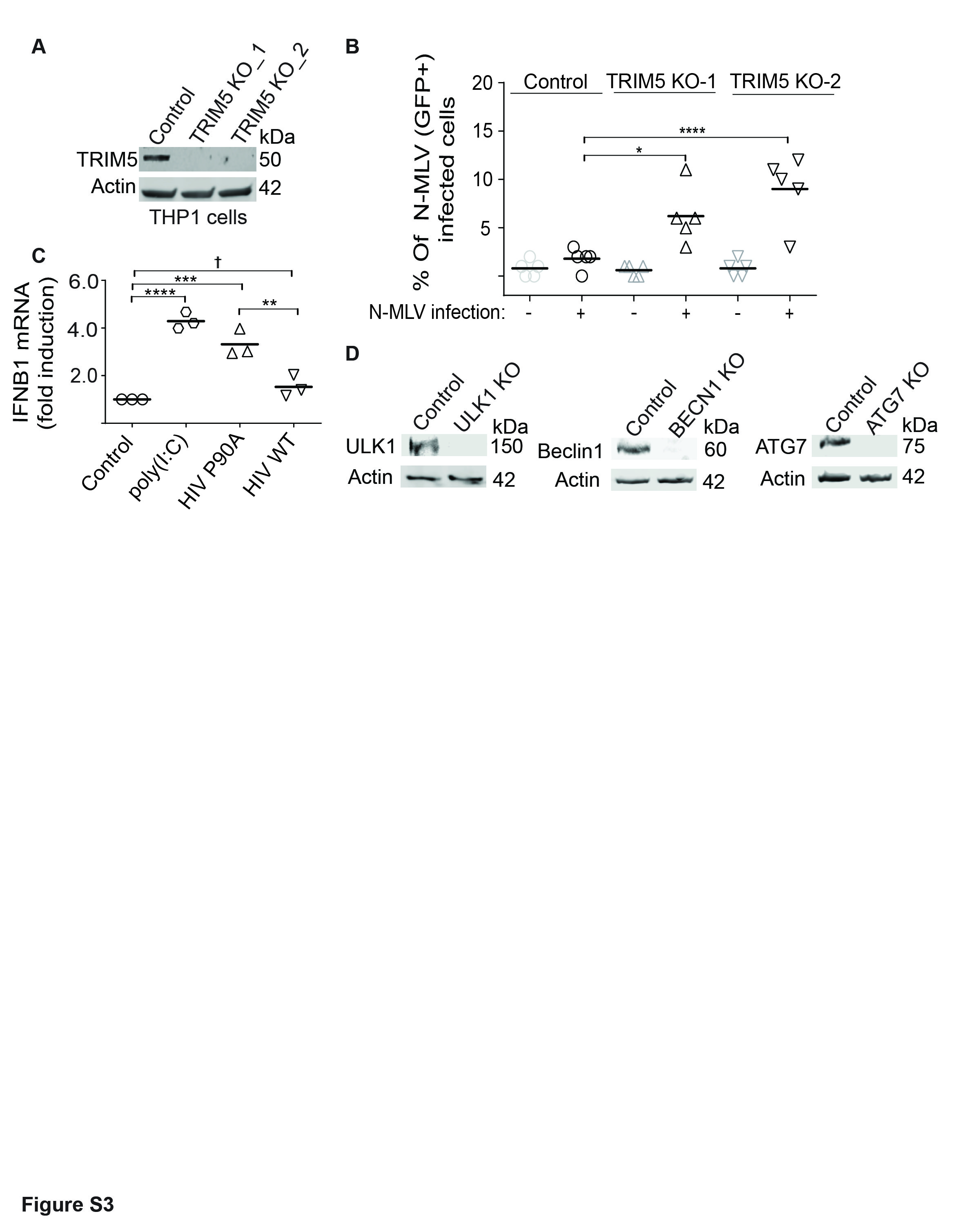

### Figure S4

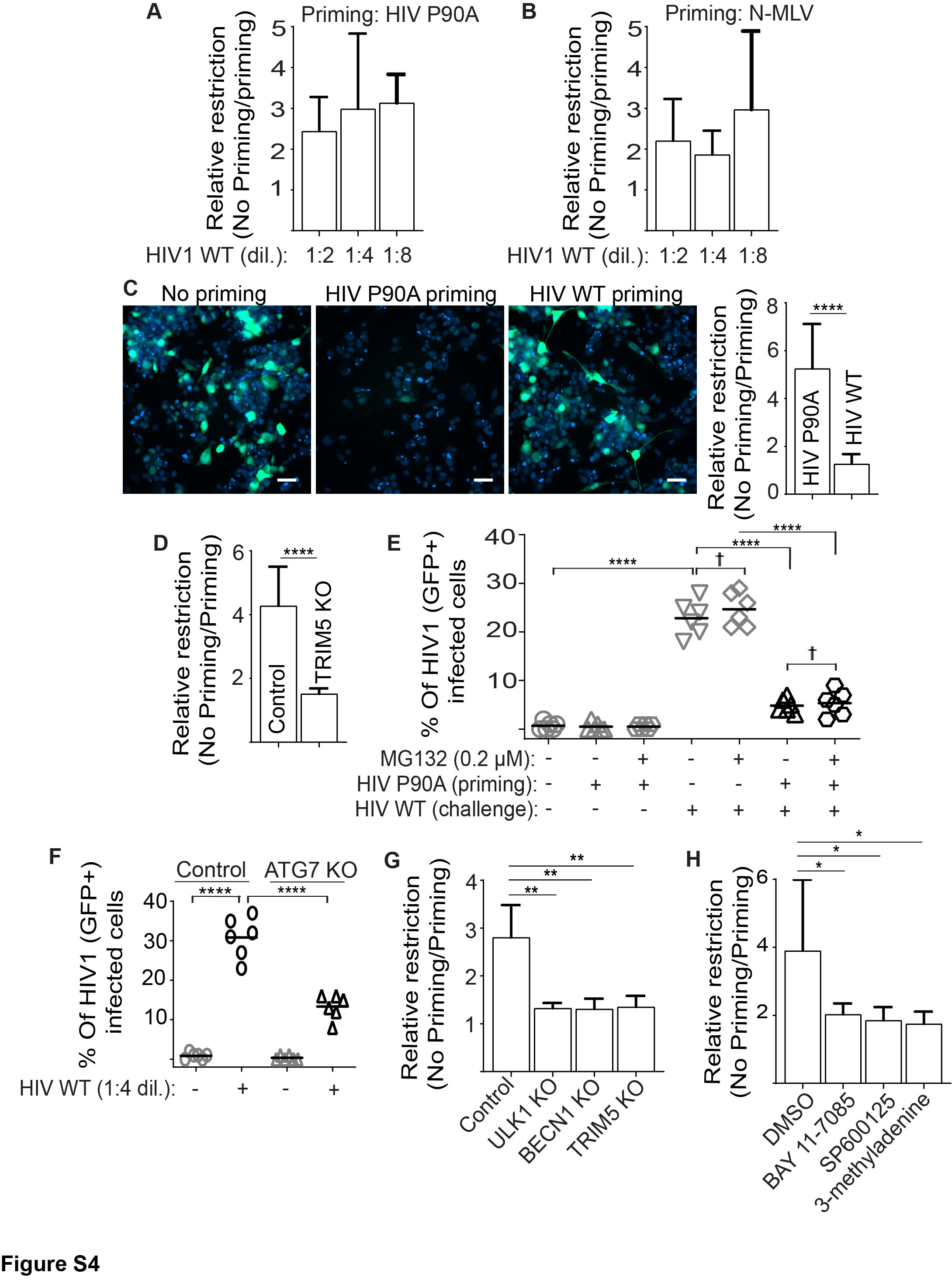

### Figure S5

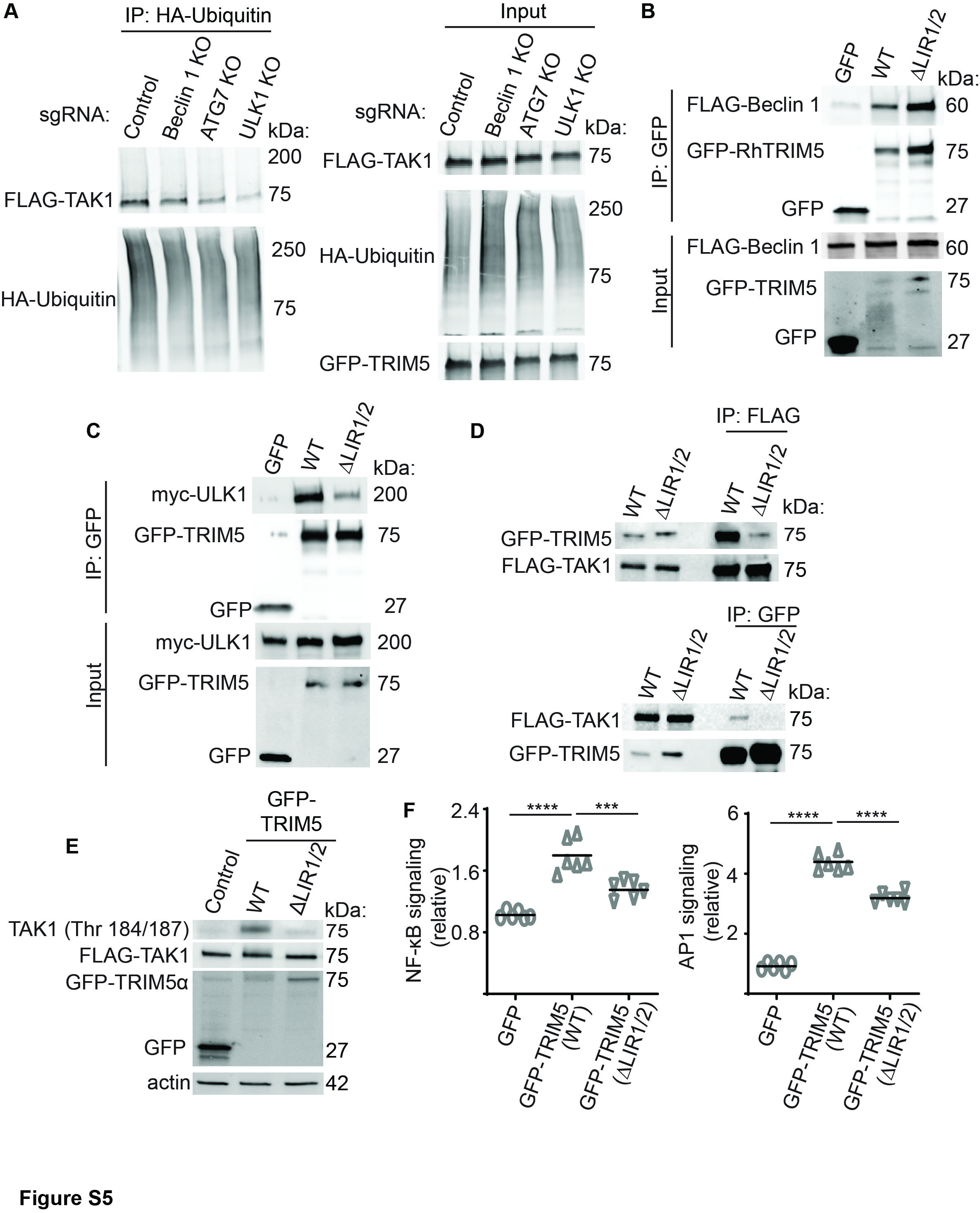
